# Experimental and field data support habitat expansion of the allopolyploid *Arabidopsis kamchatica* owing to parental legacy of heavy metal hyperaccumulation

**DOI:** 10.1101/810853

**Authors:** Timothy Paape, Reiko Akiyma, Teo Cereghetti, Yoshihiko Onda, Akira Hirao, Tanaka Kenta, Kentaro K. Shimizu

**Affiliations:** Department of Evolutionary Biology and Environmental Studies, University of Zurich, Winterthurerstrasse 190, CH-8057 Zurich, Switzerland; Sugadaira Montane Research Center, University of Tsukuba, Ueda, Nagano, Japan; Kihara Institute for Biological Research, Yokohama City University, Yokohama, Kanagawa, 244-0813, Japan

**Keywords:** polyploid speciation, adaptation, heavy metal hyperaccumulation, homeolog, expression ratio, quantitative variation

## Abstract

Little empirical evidence is available whether allopolyploid species combine or merge adaptations of parental species. The allopolyploid species *Arabidopsis kamchatica* is a natural hybrid of the diploid parents *A. halleri*, a heavy metal hyperaccumulator, and *A. lyrata*, a non-hyperaccumulating species. Zinc and cadmium were measured in native soils and leaf tissues in natural populations, and in hydroponic cultures of *A. kamchatica* and *A. halleri*. Pyrosequencing was used to estimate homeolog expression ratios. Soils from human modified sites showed significantly higher Zn concentrations than non-modified sites. Leaf samples of *A. kamchatica* collected from 40 field localities had > 1,000 µg g^-1^ Zn in over half of the populations, with significantly higher amounts of Zn concentrations in plants from human modified sites. In addition, serpentine soils were found in two populations. Most genotypes accumulated >3000 µg g^-1^ of Zn in hydroponic culture with high variability among them. Genes involved in hyperaccumulation showed a bias in the *halleri*-derived homeolog. *A. kamchatica* has retained constitutive hyperaccumulation ability inherited from *A. halleri*. Our field and experimental data provides a compelling example in which the inheritance of genetic toolkits for soil adaptations likely contributed to the habitat expansion of an allopolyploid species.

## Introduction

The ecological advantages of whole genome duplication (WGD) and its implication on species habitats has been speculated for decades (Ohno 1970; Stebbins 1971; Comai 2005; Soltis *et al*. 2014). Allopolyploids are interspecific hybrids of two or more parental species that inherit unreduced sets of chromosomes from their parental species. Allopolyploid species may have the potential to combine or merge parental adaptations (Doyle *et al*. 2008; Buggs *et al*. 2014) which may provide enhanced abilities to tolerate extreme conditions (Levin 2002; Adams 2007). Inherited adaptations to abiotic conditions may also provide allopolyploids with the ability to inhabit species distributions beyond either parental species (Blaine Marchant *et al*. 2016; Shimizu-Inatsugi *et al*. 2016; Van de Peer *et al*. 2017). The evidence for range expansion following allopolyploidization has been largely circumstantial (Hoffmann 2005; Blaine Marchant *et al*. 2016) and identifying ecologically relevant traits that may contribute to broadening habitats by allopolyploid species is essential (Ramsey & Ramsey 2014).

The allotetraploid species *Arabidopsis kamchatica* has one of the broadest distributions of any *Arabidopsis* species (Hoffmann 2005; Shimizu-Inatsugi *et al*. 2009) and offers a unique system to examine environmental responses at phenotypic and molecular levels. The species is a natural hybrid of the diploid parents *A. halleri*, a hyperaccumulator of the heavy metals cadmium (Cd) and zinc (Zn), and *A. lyrata*, a non-hyperaccumulator. Hyperaccumulating plant species such as *A. halleri,* transport large amounts of toxic heavy metals such as Cd and zinc Zn from roots to the aerial parts of the plant and are also hyper-tolerant to high concentrations of these metals in soils (Bert *et al*. 2000; Pauwels *et al*. 2006; Krämer 2010), while non-hyperaccumulators such as *A. lyrata* prevent transport of heavy metals from roots to shoots (Paape *et al*. 2016).

Metal concentrations in soils are an important factor for environmental niches. Critical toxicity of soils for plants has been defined using arbitrary thresholds, 100-300 µg g^-1^ for Zn and 6-8 µg g^-1^ for Cd (Krämer 2010). Bert *et al*. (2002) proposed similar values of soil metals, in which soil with more than 300 µg g^-1^ Zn or 2 µg g^-1^ Cd be classified as metalliferous (or metal-contaminated) soil according to the French agricultural recommendation. Stein *et al*. (2017) classified metalliferous and non-metalliferous soils based on a multifactorial metal analysis rather than using threshold values. Hyperaccumulation in plants has been defined to be > 3000 µg g^-1^ of Zn and >100 µg g^-1^ Cd in leaves (Krämer 2010). The hyperaccumulator species *A. halleri* is living in both contaminated soils near mines and non-contaminated soils, and plants from both types of soils can hyperaccumulate (Bert *et al*. 2002; Stein *et al*. 2017). Because hyperaccumulation is constitutive (species-wide) in *A. halleri*, it may have evolved hyperaccumulation as a mechanism to extract high amounts of heavy metals from metal-deficient soils for chemical defense (Boyd 2007). It has been demonstrated experimentally that *A. halleri* plants treated with Cd or Zn are more resistant to specialist and generalist insect herbivores (Kazemi-Dinan *et al*. 2014, 2015). Zinc concentration in leaves of *A. halleri* above 1,000 µg g^-1^ was shown to be effective in chemical defense. The other diploid parent, *A. lyrata* has known adaptations to serpentine soils in some regions of the species distribution but this appears to be local adaptation (Turner *et al*. 2010; Arnold *et al*. 2016) and it is not a constitutive trait. It is not known whether the habitats of *A. kamchatica* encompass human contaminated sites.

Hyperaccumulation in *A. kamchatica* appears weakened compared to the *A. halleri* parent, but is greatly increased compared to the *A. lyrata* parent. In experimental conditions, four natural genotypes of *A. kamchatica* were shown to accumulate Zn in leaf tissues to about half of the *A. halleri* parent, but 10-100 times more than the non-hyperaccumulating *A. lyrata* parent (Paape *et al*. 2016). It is not surprising that the trait distribution in these four genotypes is between the two parents, considering the divergence in hyperaccumulation between the diploid progenitors of *A. kamchatica*. However, in this small sample size, little variation in Zn accumulation among genotypes was detected in leaf tissues. While it is clear that some genotypes of *A. kamchatica* can accumulate substantial amounts of heavy metals under experimental conditions, combining data from the laboratory and *in natura* is necessary to study adaptation (Shimizu *et al*. 2011; Yamasaki *et al*. 2017). Specifically, very little is known about whether *A. kamchatica* hyperaccumulates in field conditions (Kosugi *et al*. 2016) or how much quantitative variation in hyperaccumulation exists in the species.

The genetic basis of Cd and Zn hyperaccumulation has been studied extensively in *A. halleri* using comparative transcriptomics (Filatov *et al*. 2006; Talke 2006), QTL mapping using *A. halleri* and *A. lyrata* parents (Courbot *et al*. 2007; Willems *et al*. 2007; Frérot *et al*. 2010), and functional genetics (Hanikenne *et al*. 2008). These studies have revealed important metal transporter genes involved in the uptake, root to shoot transport, and cellular detoxification of heavy metals that showed enhanced expression in *A. halleri* compared to *A. thaliana* or *A. lyrata* orthologs (Hanikenne & Nouet 2011). In *A. kamchatica*, these genes are inherited homeologs of these genes from both of the diploid progenitors. Relative expression levels of both homeologs may exhibit similar levels as in the parental species (“parental legacy”) resulting in expression bias (Buggs *et al*. 2014; Yoo *et al*. 2014) in genes involved in hyperaccumulation that were inherited from *A. halleri*. Genome wide transcriptomics analysis of two *A. kamchatica* genotypes showed that expression ratios of many heavy metal transporters were strongly biased toward the *A. halleri* derived homeolog (Paape *et al*. 2016), as expected if *A. kamchatica* inherited hyperaccumulation ability from *A. halleri*. If this bias in important heavy metal transporters is detected universally among widespread genotypes of *A. kamchatica*, it would suggest that parental legacy or constitutive expression has been maintained throughout the species distribution.

The two main loci involved in hyperaccumulation in *A. halleri* are *HMA4 (HEAVY METAL ATPASE4*), which is an ATPase transporter protein, and *MTP1* (*METAL TOLERANCE PROTEIN1* also called *ATCDF1* or *ZAT1*), a cation diffusion facilitator (CDF) protein (Frérot *et al*. 2010). In *A. halleri*, *HMA4* and *MTP1* are constitutively highly expressed even under low metal conditions (Talke 2006; Paape *et al*. 2016) and both have known paralogous duplications in *A. halleri* (but single copy in other *Arabidopsis* species, Briskine *et al*. 2017), together resulting in higher expression than non-hyperaccumulating congeners (Hanikenne et al. 2008; Shahzad et al. 2010). Another ATPase, *HMA3,* involved in vacuolar sequestration of Zn (Morel *et al*. 2008), is also constitutively expressed in *A. halleri*. The gene *MTP3* (same protein family as *MTP1*) is mainly expressed in root tissues and is involved in vacuolar sequestration of Zn in roots of *A. thaliana* (Arrivault *et al*. 2006). By sequestering Zn in the roots, it is thought that *MTP3* prevents toxic heavy metal transport from roots to shoots by retaining Zn in the roots in the non-hyperaccumulating species *A. thaliana*. The zip transporter *IRT3* and natural resistance associated macrophage protein *NRAMP3* encode iron (Fe) transporters that are co-regulated by Fe and Zn, and show significant expression differences between the hyperaccumulator *A. halleri* and non-accumulators *A. thaliana* (Talke 2006; Lin *et al*. 2009; Shanmugam *et al*. 2011) and *A. lyrata* (Filatov *et al*. 2006; Paape *et al*. 2016).

We propose that heavy metal hyperaccumulation in *A. kamchatica* is an ecologically-relevant trait that may broaden the habitat in an allopolyploid species. However, we know very little about whether plants hyperaccumulate in nature or how much variation in hyperaccumulation exists in the species. In this study, we set out to answer the following questions. (1) Do the habitats of *A. kamchatica* contain heavy metals and is there evidence of naturally- and artificially-generated metalliferous soils? (2) Does hyperaccumulation of Cd and Zn occur in natural populations of *A. kamchatica*? (3) Is there quantitative variation in leaf and root accumulation in *A. kamchatica*, and are there genotypes that can hyperaccumulate as much as *A. halleri*? (4) Is *A. halleri* a more efficient hyperaccumulator than *A. kamchatica* at all concentrations of Zn treatments? (5) Do genes involved in heavy metal transport and detoxification show a bias in expression ratios of homeologs among *A. kamchatica* genotypes that may be derived from the hyperaccumulating parent, *A. halleri*? By combining field data with phenotyping in experimental conditions and homeolog specific gene expression estimates, we were able to provide the first range-wide study of constitutive heavy metal hyperaccumulation in an allopolyploid species.

## Results

### Zn accumulation in soils and leaf tissues in wild populations

*Arabidopsis kamchatica* is a ruderal species which grows both in human–modified and natural habitats (see definitions in next paragraph) but none of the samples in this study were found where mining activities occurred. We sampled leaf tissues from 40 populations of *A. kamchatica* from Japan and Alaska, USA to quantify Zn. We obtained soils of 37 of these localities and measured Zn concentration of the soils. The concentration of Zn in soils among the sites ranged from 18.7 to 642.8 (average 150 µg g^-1^, Fig. 1, Table S1-S3). The Zn concentrations in soils of 18 sites were above 100 µg g^-1^, and five sites were above 300 µg g^-1^. This indicated that a considerable number of sites where *A. kamchatica* is growing is are above the critical toxicity of 100-300 µg g^-1^ for Zn, while others are below the thresholds.

**Figure 1.**
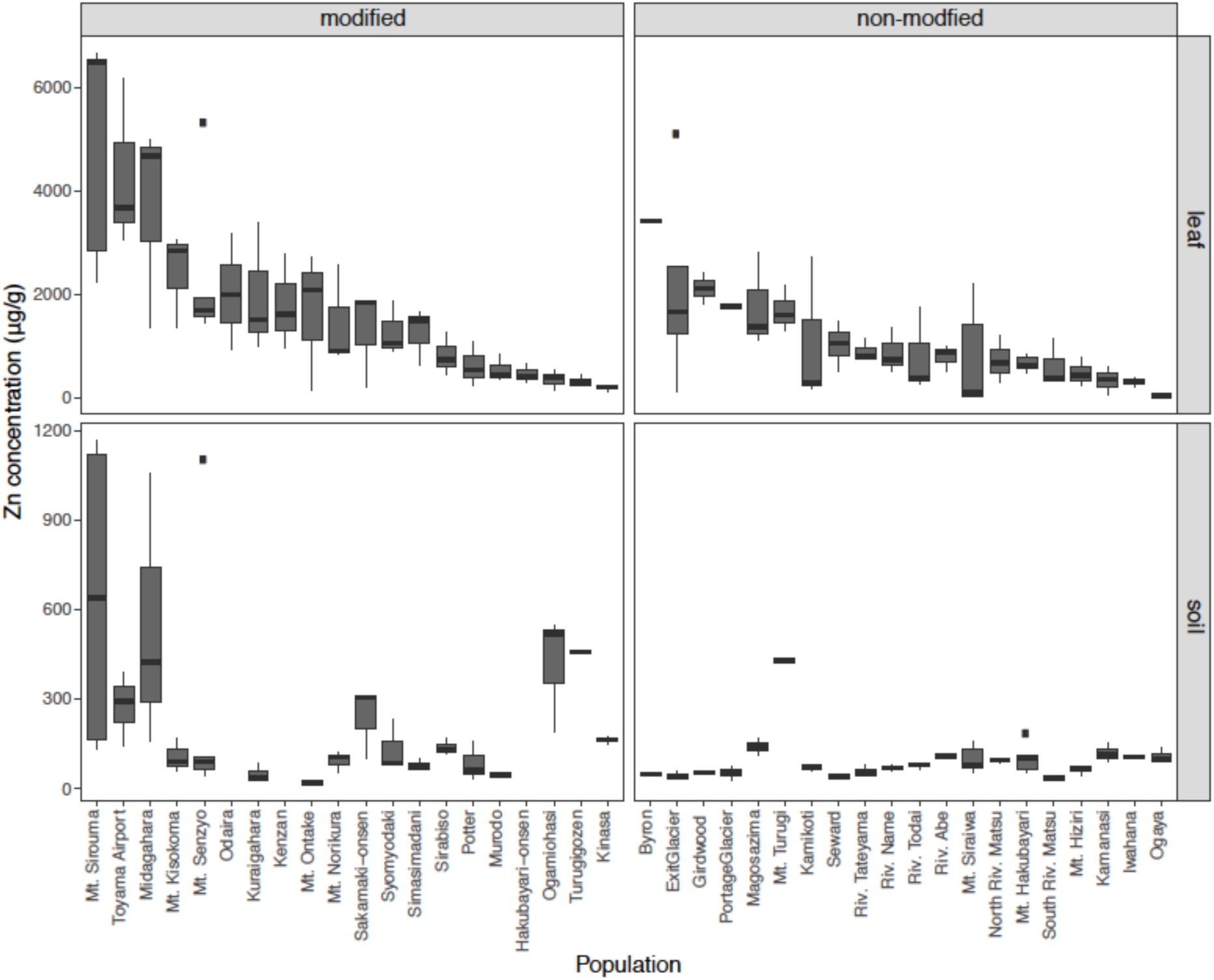
Leaf concentrations of zinc (Zn) from *A. kamchatica* plants collected in 40 natural populations of (upper panel). Soil concentrations of Zn (lower panel) at 37 corresponding sites. Populations were grouped into modified or non-modified. The populations are sorted from highest to lowest Zn in the leaf samples. The y-axis is Zn concentration in µg per gram of dry weight. Note different y-axis scales for leaf and soil. Boxplots show center line: median; box limits: upper and lower quartiles; whiskers: 1.5 × interquartile range; points: outliers. The correlations between leaf accumulation and soil concentrations: (modified) Pearson’s *r* = 0.68, (non-modified) Pearson’s *r* = 0.007, see also Figure S1. See Table S1 for mean values, variance, number of replicates, and location information.

Observations made during plant and soil collection were that several of the sites had some evidence of human modification such as buildings, mountain lodges, fences or paved roads. To examine whether *A. kamchatica* could adapt to naturally- and artificially- generated metalliferous soils, we classified the localities into human-modified or non-modified sites. In human–modified sites, there was artificial construction (buildings and fences on concrete base; and paved roads and riverside) close (< 10 m) to the *A. kamchatica* plants that were collected (Table S4). The second category encompasses the sites without artificial construction. The average Zn concentration at modified sites was significantly higher than the non-modified sites (p = 0.0005, Table S3, Fig. 1), suggesting the existence of artificially–introduced Zn. Most of the sites with > 300 µg g^-1^ Zn in soils are human-modified sites (i.e. Mt. Sirouma, Midagahara, Turugigozen, Ogamiohasi; Table S4). Yet, many sites without human modification also showed considerable amounts of Zn (seven sites with >100 µg g^-1^ Zn, one of them >300 µg g^-1^ Zn), and are likely to reflect natural geology (Schlüchter *et al*. 1981). These data suggest that the habitat of *A. kamchatica* encompasses localities with high Zn in soil, both by natural geology and human modification, as well as sites with low Zn in either category.

Next, we measured the Zn concentration in leaf tissues of *A. kamchatica* collected at each of the 40 populations. The average accumulation of Zn was 1,416 µg g^-1^ dry weight (DW) among 126 plants from the 40 populations. We detected > 1,000 µg g^-1^ of Zn in 21 of the 40 populations and > 3,000 µg g^-1^ in four populations representing both modified and non-modified sites (Figure 1, Table S2). The highest concentration of Zn was found in three plants at the Mt. Sirouma site (6,500 – 6,661 µg g^-1^) in Japan (2835 meters above sea level), a human modified site with the highest level of Zn in the soils (Fig. 1). We found that plants from human-modified habitats had on average higher quantities of Zn (mean Zn concentration 1,815 µg g^-1^) than non-modified sites (mean Zn concentration 1,004 µg g^-1^) and the difference was significant (p = 0.00136, Fig. 1, Tables S1-S3). In human-modified habitats there was a significant positive correlation between leaf and soil concentrations of Zn (*r* = 0.68, p < 10^-5^) suggesting that the increased concentrations of Zn in the soils at these sites increased the availability of this metal for uptake by plants. By contrast, there was no correlation between leaf and soil concentrations of Zn in non-modified habitats (*r* = 0.007, p = 0.95; Fig. S1). We found that *A. kamchatica* accumulated high levels of Zn in the field and when there is greater availability of Zn in the soils, plants tend to have higher levels of Zn in the leaves as we found in many of the modified sites.

In the parental species *A. halleri*, collected from two sites in Russia and two sites in Japan, we detected > 3,000 µg g^-1^ of Zn in leaves in all four populations, with the highest levels found in plants from the Tada mine site in Japan (average of six replicates = 16,068 µg g^-1^ (Table S1)). Soils were collected at the two mine sites in Japan (Tada and Omoidegawa (OMD)) (Table S4), and the Zn concentration (1,317 – 2,490 µg g^-1^ Zn) was several times higher than any other sites with *A. kamchatica*.

### Cadmium and serpentine soils in the habitat of A. kamchatica

Because *A. halleri* is also a known hyperaccumulator of Cd, we measured the concentration of Cd in soils and in leaf tissues of *A. kamchatica*. The soils from all measured sites of *A. kamchatica* had < 2 µg g^-1^ Cd, indicating that soils are not contaminated with this heavy metal in its habitats. The average accumulation of Cd in leaves among the populations of *A. kamchatica* was 1.8 µg g^-1^ (Figure 2, Table S3). The Cd concentration in leaves in the five populations with the highest Cd ranged from 3.6 to 8.9 µg g^-1^, while all other populations had less than 3 µg g^-1^ of Cd in leaves (Table S1). The maximum amount of Cd in any single plant was 12.1 µg g^-1^ (Table S2, S3) and was found at the Mt. Siraiwa site, which also had the highest Cd levels in soils. However, this level of Cd accumulation in leaves is much lower than the threshold defined for Cd hyperaccumulation (>100 µg g^-1^ Cd) in leaves (Krämer 2010). Unlike for Zn, there was no significant difference in Cd concentrations in the soils or leaf tissues between the modified or non-modified sites (Table S3). Moreover, there was no correlation between leaf and soil concentrations of Cd in the modified habitats, but there was a positive correlation in the non-modified habitats (Fig. S1). We also found very little correlation between Cd and Zn leaf accumulation (*r* = 0.08, p = 0.39), likely due to much less overall Cd in soils and leaves compared with Zn. In summary, leaf and soil concentrations of Cd are below critical toxicity levels for plants (Krämer 2010) and are negligible compared to Zn concentrations in the same populations.

**Figure 2.**
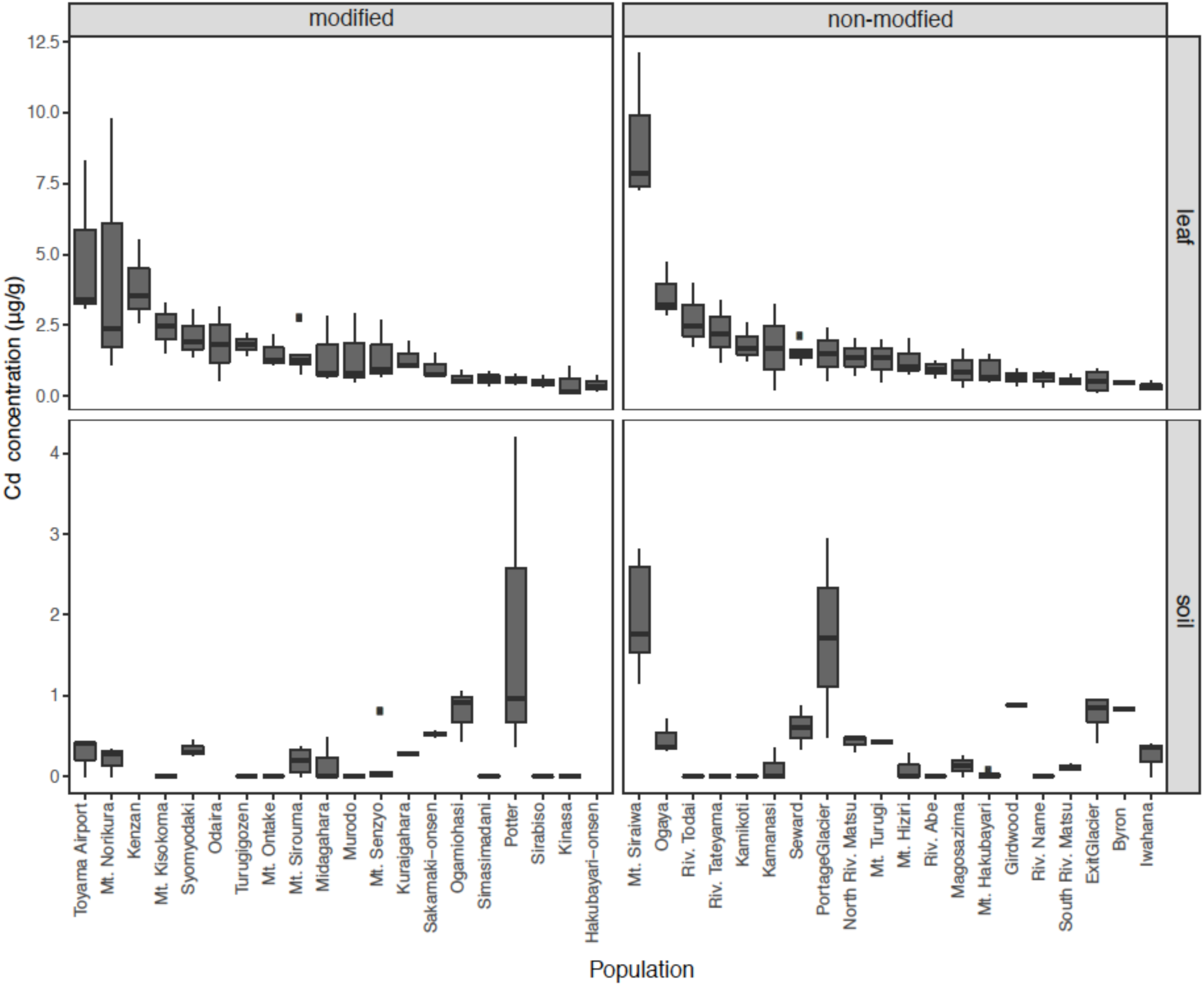
Leaf concentrations of cadmium (Cd) from *A. kamchatica* plants collected in 40 natural populations of (upper panel). Soil concentrations of Cd (lower panel) at 37 corresponding sites. Populations were grouped into modified or non-modified. The populations are sorted from highest to lowest Zn in the leaf samples. The y-axis is Cd concentration in µg per gram of dry weight. Note different y-axis scales for leaf and soil. Boxplots show center line: median; box limits: upper and lower quartiles; whiskers: 1.5 × interquartile range; points: outliers. The correlations between leaf accumulation and soil concentrations: (modified) Pearson’s *r* = -0.08, (non-modified) Pearson’s *r* = 0.58, see Figure S1. See Table S1 for mean values, variance, number of replicates, and location information.

For *A. halleri*, soil at the two Japanese mine populations contained 5.0 – 6.9 µg g^-1^ of Cd (Table S4), which is comparable to the critical toxicity defined by Krämer (2010) (6-8 µg g^-1^). Leaf Cd concentration of the two Japanese and two Russian populations were measured, and one of them was above the >100 µg g^-1^ threshold of hyperaccumulation (1,267 µg g^-1^at Tada mine). This supports a previous report on facultative hyperaccumulation of Cd by *A. halleri* (Stein et al. 2017). While our main objective of the soil and leaf sampling of *A. kamchatica* was to quantify the heavy metals Cd and Zn, we also found high levels of magnesium (Mg) and nickel (Ni) in the soils of two populations in Japan (Mt. Sirouma and Mt. Hakubayari) (Figure S2, Figure S3; Table S4). These two sites had nearly an order of magnitude greater Mg and Ni than all other populations (the Mt. Sirouma site also contained the highest soil concentration of Zn of all of the *A. kamchatica* sites (643 µg g^-1^)). High concentrations of Mg and Ni, and low calcium to magnesium ratios (Ca:Mg) indicate serpentine soils (Kazakou *et al*. 2008). Consistent with this, the Ca:Mg in soils Mt. Sirouma (Ca:Mg = 3.71e^-05^) and Mt. Hakubayari (Ca:Mg = 3.45e^-05^) were at least an order of magnitude lower than the average among all other Japanese populations which was 1.60e^-03^ (sd. 2.60e^-03^). Therefore, the high levels of Mg and Ni, in addition to low Ca:Mg, indicates that *A. kamchatica* is living on serpentine soils, and the mountain range including Mr. Sirouima and Mt. Hakubayari are known to have serpentine soils (Hatano and Matsuzawa 2008). Furthermore, the River Matu is originated in this mountain range, and the North and South River Matu sites had three times higher Ni than most populations.

### Variation of zinc accumulation in experimental conditions

To examine the genetic variation of Zn hyperaccumulation in a common environment, Zn accumulation in leaves and roots were quantified using hydroponic solution in growth chambers. We used a treatment condition of 500 µM of Zn exposure for 7 days. We examined *A. kamchatica* genotypes sampled from 19 natural populations that span the species range (Table S5a), plus one synthesized allopolyploid (20 genotypes in total). A minimum of five replicates of each genotype were measured. Linear models detected significant quantitative variation in Zn accumulation, even when outlier genotypes were removed (Table S5b). Broad sense heritability (*H*^2^) for leaf accumulation of Zn in *A. kamchatica* was estimated to be as high as 0.70, which reflects significant variation among genotypes (Figure 3, Table S6). Among the 20 plant genotypes, the average Zn accumulation in leaves was 4,562 µg g^-1^ and ranged from 1,845 – 16,213 µg g^-1^ in mean values among replicates in all genotypes (Figure 3, Table S5a). The synthesized allopolyploid used in this experiment had among the highest level of Zn accumulation in leaves, 5,845 µg g^-1^ demonstrating that high Zn accumulation can be retained following early hybridization between *A. halleri* and *A. lyrata*. To check whether Zn accumulation among these samples could be explained by native soil concentrations of Zn, we compared 10 populations of *A. kamchatica* for which both soil Zn concentration and leaf accumulation in the growth chamber experiment were available. The variance of Zn concentration in leaf tissues explained by Zn concentration in soil was not significant (p = 0.8) and could be explained mostly by plant genotype (p < 2e^-16^).

**Figure 3.**
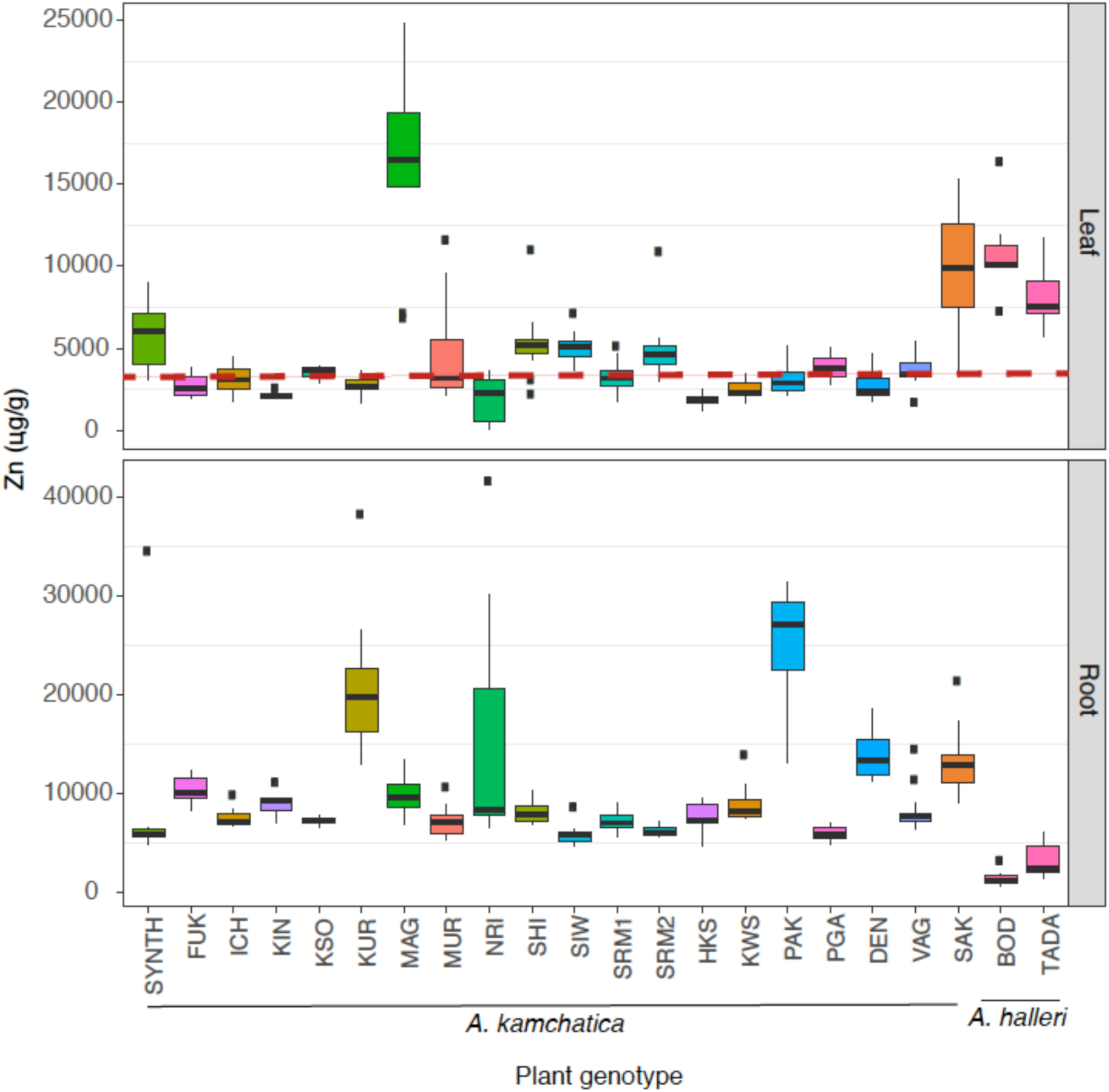
Zinc accumulation in leaf tissue (upper panel) and root tissue (lower panel) in 20 *A. kamchatica* and 2 *A. halleri* genotypes after one week of 500 µM zinc treatment in hydroponic chambers. The dashed red line is 3,000 µg, the threshold defining hyperaccumulation (Krämer 2010). Boxplots show center line: median; box limits: upper and lower quartiles; whiskers: 1.5 × interquartile range; points: outliers. See Table S5 for mean and median values, standard deviations, and number of replicates.

For comparison, we included two *A. halleri* genotypes in the analysis. The TADA and BOD genotypes accumulated 8,036 µg g^-1^ and 10,891 µg g^-1^ of Zn in leaf tissues, respectively (Figure 3, Table S5). The TADA genotype of *A. halleri* which was collected from a highly metal contaminated site (metalliferous) near the Tada mine (Briskine *et al*. 2017) showed less Zn accumulation in leaves than the BOD genotype, which was collected from a region with no mine activity (non-metalliferous site). This is consistent with other studies that demonstrated *A. halleri* from non-metalliferous sites often accumulate more than those from metalliferous sites (Bert *et al*. 2002; Stein *et al*. 2017). While the average Zn accumulation in leaves of *A. kamchatica* is about half compared with *A. halleri*, two natural genotypes had high Zn accumulation in leaves at similar or higher levels compared to *A. halleri*. The SAK genotype from Sakhalin Island had mean Zn accumulation of 9,562 µg g^-1^ in leaf tissues among 12 replicates, with a maximum of 15,348 µg g^-1^ in one individual. The MAG genotype (from Magosazima, Japan) had mean Zn accumulation of 16,213 µg g^-1^ in leaf tissues among 8 replicates and a maximum of 24,809 µg g^-1^ in one individual (Table S5).

A major difference between *A. kamchatica* and *A. halleri* was in the level of Zn accumulation in the roots. Accumulation of Zn in the roots of the 20 *A. kamchatica* genotypes was between 5,980 and 25,650 µg g^-1^, whereas in *A. halleri,* Zn concentrations in roots were 3,190 µg g^-1^ for the TADA genotype and 1,525 µg g^-1^ for the BOD genotype, lower than all *A. kamchatica* genotypes (Figure 3, Table S5a). The shoot to root ratio of Zn accumulation was greater than one (i.e., higher Zn concentrations in leaves than roots) for both genotypes of *A. halleri*, while all but one of the *A. kamchatica* genotypes had a leaf to root ratio in Zn accumulation greater than one (the MAG genotype had a leaf to root ratio greater than one (Table S5a)). There was no significant correlation between leaf and root accumulation in *A. kamchatica* (*r* = -0.06, p = 0.37), indicating that genotypes with high metal accumulation in leaves do not necessarily have lower accumulation in roots, and vice versa. Together, these comparisons suggest that leaf tissues, rather than roots, are a sink for heavy metals in *A. halleri,* and that the diploid species has a more efficient mechanism of root to shoot transport of Zn than *A. kamchatica*.

We also included a synthesized allopolyploid generated by a cross between Asian *A. halleri* ssp. *gemmifera* (TADA genotype, also used in this experiment) and Siberian *A. lyrata* ssp. *petrea* (Akama *et al*. 2014; Paape *et al*. 2018). The synthetic polyploid showed a higher level of Zn accumulation in leaves than most natural genotypes but it was significantly lower than the *A. halleri* parental genotype (TADA) (p = 0.04, Table S6). In contrast to natural *A. kamchatica* that experienced evolutionary changes after polyploid speciation, the synthesized allopolyploid provides direct experimental evidence that Zn hyperaccumulation in *A. halleri* can be inherited in *A. kamchatica,* but that the trait is attenuated due to genome merging with the non-hyperaccumulator *A. lyrata*.

Next, we wanted to test whether plant genotypes responded similarly when the Zn treatment was increased from 500 µM to 1,000 µM Zn. Using nine of the twenty *A. kamchatica* genotypes that were used in the previous (500 µM) experiment, this time treated with 1,000 µM Zn, we found significant increases in Zn accumulation in the leaves in all but two genotypes (Figure S4). The two genotypes (MAG and SAK) that showed the highest Zn accumulation in the 500 µM treatment, did not significantly increase Zn accumulation in leaves when treated with 1,000 µM of Zn. This demonstrated that Zn accumulation in *A. kamchatica* leaf tissues may be increased by exposure to higher heavy metals in the roots, but for the two highest accumulating genotypes, it may suggest a maximum threshold or saturation of heavy metals in the leaves under high Zn dosage.

### Short-term Zn transport comparisons between A. halleri and A. kamchatica

Based on the hydroponic experiments above, all *A. kamchatica* genotypes were able to accumulate substantial amounts of Zn in the leaf tissues and none would be considered non-accumulating genotypes that more closely resemble *A. lyrata* (based on previous studies using *A. lyrata*, Filatov *et al*. 2006; Willems *et al*. 2007; Paape *et al*. 2016). We also know that both *A. halleri* and *A. kamchatica* can accumulate large amounts of Zn even when exposed to relatively low concentrations of Zn in the soil, but in our experiments above, *A. halleri* had a higher ratio of leaf to root Zn accumulation than nearly all *A. kamchatica* genotypes. We therefore compared the relative efficiency of Zn transport between *A. halleri* and *A. kamchatica* over 10-fold gradients of Zn treatment for 48 hours, so that a rapid transport from roots to leaves upon exposure to various concentrations of Zn may be detected. In this experiment, we tested Zn accumulation in leaves and roots of one *A. halleri* genotype and three *A. kamchatica* genotypes. The three *A. kamchatica* genotypes were selected to represent different geographic regions: Alaska (PAK), Japan (MUR), and Sakhalin Island (SAK) and the *A. halleri* genotype (TADA) comes from the Tada mine area in Japan.

In leaf tissues, *A. halleri* accumulated > 2,000 µg g^-1^ when treated with only 10 µM of Zn. This was a four-fold increase compared with the 1 µM treatment condition. There was no significant difference in Zn accumulation in leaves of *A. halleri* between the 10 and 100 µM treatments, then nearly 2-fold increase between the 100 and 1000 µM treatments (Figure 4A). By contrast, *A. kamchatica* showed the largest fold-increase in Zn accumulation in leaf tissues following the largest Zn treatment dosage of 1,000 µM. Following each of the three treatment conditions (10, 100, and 1,000 µM), *A. halleri* accumulated significantly more Zn in leaves than *A. kamchatica*, except for one *A. kamchatica* genotype (PAK) at the 1,000 µM treatment. It was not until the highest Zn treatment (1,000 µM) that *A. kamchatica* accumulated the same amount (2,000 µg g^-1^) of Zn as did *A. halleri* at only 10 µM of Zn treatment.

**Figure 4.**
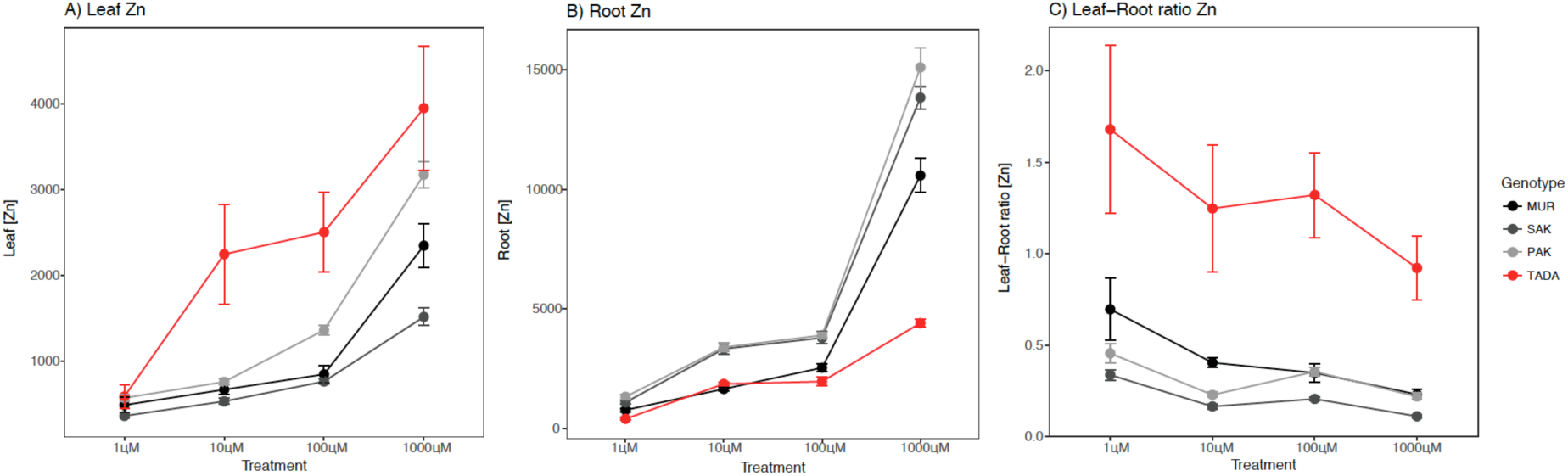
Leaf (A) and root (B) accumulation of Zn by *A. halleri* (TADA, red), and three *A. kamchatica* genotypes (PAK (light gray), SAK (dark gray), MUR (black)) at four Zn treatment conditions: 1 µM, 10 µM, 100 µM, and 1000 µM. (C) The ratio of leaf to root Zn accumulation at each treatment condition.

In root tissues, *A. halleri* showed little increase in Zn accumulation until the 1,000 µM treatment, and Zn accumulation in roots did not exceed 5,000 µg g^-1^ in any treatment condition (Figure 4B). The shoot to root ratio of Zn concentrations in each of the three treatment conditions was ≥ 1 for *A. halleri* (Figure 4C). For *A. kamchatica,* Zn concentrations in roots were significantly greater than *A. halleri* in the three treatment conditions (with the exception of the MUR genotype at the 10 µM treatment). The 1,000 µM treatment resulted in the largest Zn accumulation in the roots, ≥ 10,000 µg g^-1^ for all three *A. kamchatica* genotypes (Figure 4B), which was twice as high as *A. halleri* at this treatment. The high Zn accumulation in the roots of *A. kamchatica* resulted in shoot to root ratios ≤ 0.5 at all treatments, less than *A. halleri* at each Zn treatment condition (Figure 4C). These results further show that *A. halleri* is a more efficient hyperaccumulator of Zn, especially when exposed to low concentrations.

### Homeolog expression ratios of candidate genes

We used pyrosequencing to quantify homeolog expression ratios for six known candidate genes involved in heavy metal tolerance and hyperaccumulation in the leaves and roots of 15 genotypes of *A. kamchatica.* Expression ratios were quantified before and after Zn treatments. The genes were selected based on previous studies that estimated expression differences between diploid congeners (Becher *et al*. 2004; Filatov *et al*. 2006; Talke 2006) and functional assays in plants and/or yeast (Table S8). We found an overall trend for *A. halleri-*derived (H-origin) homeologs to be expressed at a higher ratio compared to the *A. lyrata*-derived (L-origin) homeologs for all of the genes tested (Figure 5), although considerable variation was observed. Linear models showed that gene, plant genotype, and tissue, all contributed to a significant proportion of the variance in expression ratios (p-values: < 2.2e^-16^, 3.52e^-13^, and 1.02e^-10^ respectively, Table S9). The genes *IRT3*, *MTP3*, and *NRAMP3* all showed significant variation among genotypes (p-values: 7.15e^-07^, 3.89e^-14^, and 1.72 e^-11^ respectively), while the genes *HMA3*, *HMA4* and *MTP1* showed no significant among-genotype variation in expression ratios (p-values: 0.72, 0.72, and 0.21 respectively).

**Figure 5.**
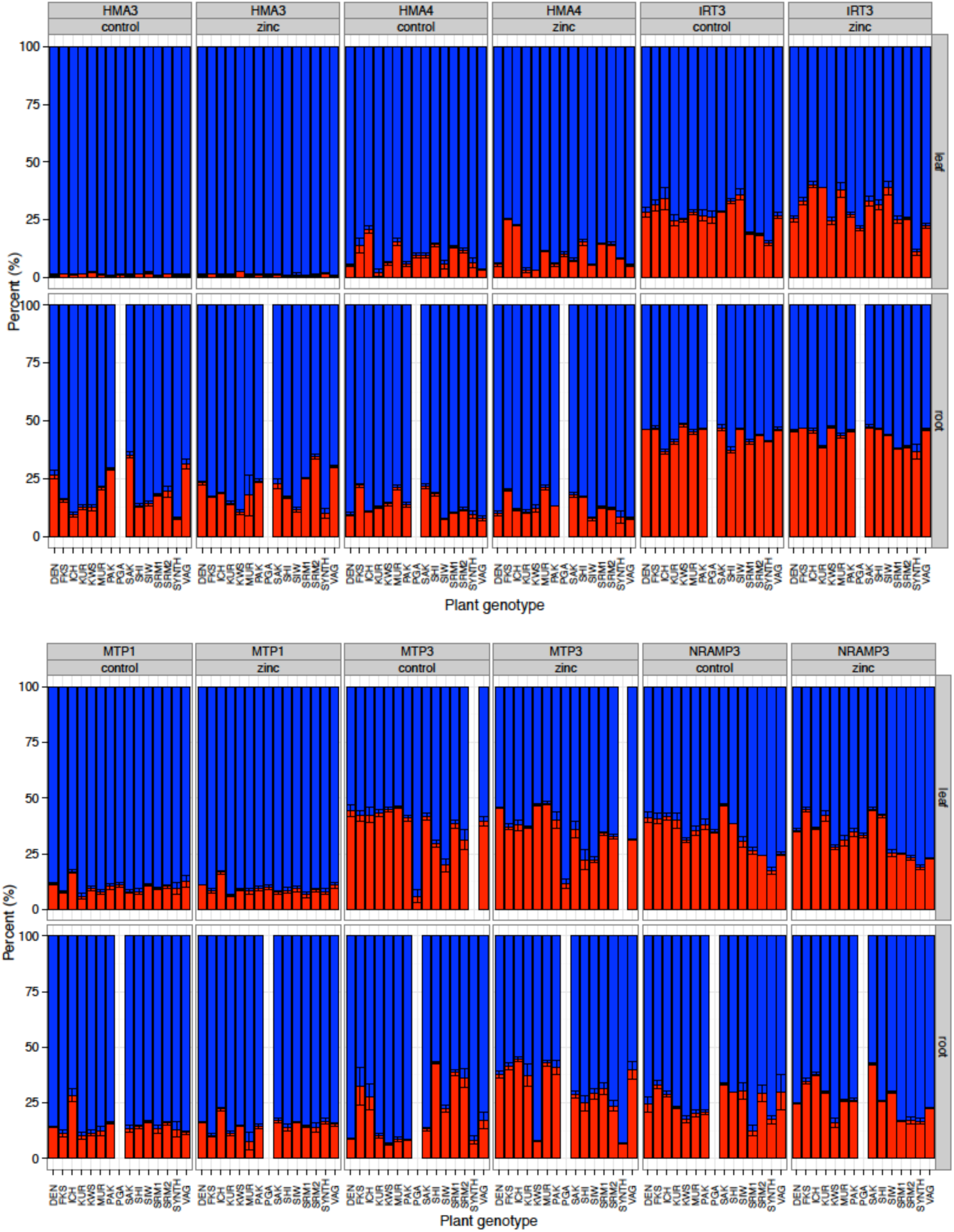
Ratios of homeolog expression estimated by pyrosequencing for 6 genes in 15 *A. kamchatica* genotypes (14 for root tissue) under control conditions (1µM of Zn) and treated (500 µM of Zn) sampled after 48 hrs. Red bars represent the *A. lyrata* derived homeolog expression, and blue bars are the *A. halleri* derived homeolog expression. Each bar represents an *A. kamchatica* genotype. Root data for the PGA genotype was not collected.

Zinc treatment had no significant effect on expression ratios when compared with control conditions when both tissues were included in the model (treatment p-value = 0.64), and when tissues were analyzed separately (leaf tissue only, p = 0.66; root only, p = 0.12). This reflects the constitutive expression of the H-origin metal transporters. Furthermore, we found no significant relationship between homeolog expression ratios and Zn hyperaccumulation levels in our experiments: the model coefficient for the interaction between gene and treatment was non-significant (gene x treatment interaction p = 0.18), and the coefficients for each individual gene and treatment interaction were non-significant (Table S9). The one exceptional coefficient was found in the *MTP3* gene x treatment x tissue (root) which was highly significant (p = 0.0005). Whether this ratio change is due to significant up-regulation of the L-origin copy, or down-regulation in the H-origin copy remains to be tested, but the pattern is consistent with up-regulation of the L-origin copy in a previous RNA-seq experiment (Paape *et al*. 2016).

## Discussion

### Natural populations of Arabidopsis kamchatica show substantial accumulation of Zn in leaves

In the first species-wide survey of *A. kamchatica*, ion measurements in leaf samples collected from natural conditions showed that the species can accumulate large amounts of heavy metals. In particular, Zn accumulation was high in many populations despite generally low heavy metal concentrations in soils for the majority of sites. Substantial concentrations of Zn were found in *A. kamchatica* plants in more than half of the populations (> 1,000 µg g^-1^), and plants from three populations that had > 3,000 µg g^-1^ of Zn, which exceeds the threshold defining Zn hyperaccumulation for plants (Krämer 2010). Because the concentrations ranged in more than two orders of magnitudes (from 30 to > 6,000 µg g^-1^) with no clear split into two classes of hyperaccumulating and non-hyperaccumulating genotypes (Table S2), this suggests that the classification into these two categories may be too simple to study intraspecific variation of hyperaccumulation. It is noteworthy that previous experiments have demonstrated that *A. halleri* plants that accumulated > 1,000 µg g^-1^ experienced a significant reduction in herbivory compared with plants with lower Zn accumulation (Kazemi-Dinan *et al*. 2014, 2015). Therefore, a similar level of Zn in leaves could also be sufficient for *A. kamchatica* to deter insects.

We did not find plants growing in sites with obvious contamination from mining as with *A. halleri*. The influence of the *A. lyrata* genome may have reduced the ability of *A. kamchatica* to inhabit highly toxic mine sites similar to *A. halleri,* such as places like the Tada mine area. However, we did find that nearly half of the sites that we collected samples from were clearly modified by human activities, and many of them contained high concentrations of Zn in the soils. This is supported by significantly higher levels of Zn concentrations in the leaves at the modified vs. non-modified sites, as well as the strong positive correlation between soil concentration and leaf accumulation of Zn in modified sites. The use of corrugated galvanized irons and grating may have artificially increased Zn in the surrounding soils near mountain lodges and roads. It is possible that the tolerance to high levels of heavy metals inherited from *A. halleri* pre-adapted *A. kamchatica* to expand its habitat into human modified sites that contain elevated Zn in soils.

Compared with Zn, the levels of Cd in soils or in field collected *A. kamchatica* are unremarkable and below toxic thresholds. There appeared to be no anthropogenic influence on the levels of Cd in the soils, as quantities at both modified and non-modified site types only differed slightly, and not significantly. Pollution from mining activities is the most common source of high amounts of Cd in soils and plants (Alloway & Steinnes 1999), and elevated Cd from human construction is not a likely source of increased Cd as it may be for Zn. The correlation between Cd and Zn concentrations in leaf tissues in *A. kamchatica* plants was low, likely due to less exposure of plants to Cd even in the populations with higher Zn concentrations in the soils.

Most of the habitat soils of *A. kamchatica* can be considered non-metalliferous based on Bert *et al*. (2002). This is a major difference compared with *A. halleri* where many populations are known to be highly contaminated due to mining activities (Bert *et al*. 2002; Pauwels *et al*. 2006; Hanikenne *et al*. 2013; Briskine *et al*. 2017), including the two sites containing *A. halleri* in Japan that we collected samples for this study. Previous studies of European populations of *A. halleri* demonstrated that plants from both metalliferous and non-metalliferous soils can hyperaccumulate (Bert *et al*. 2000), and plants from non-metalliferous sites showed ranges of Zn accumulation in leaves between 1806 – 10,876 µg g^-1^ and Cd between 3 – 89 µg g^-1^ (Bert *et al*. 2002). The range of concentrations of Zn in leaves and soils reported in Bert *et al*. (2002) were similar to that in our collection sites containing *A. kamchatica* (Table S3), but we found a lower mean and narrower range of Cd concentrations in leaves and soils. More recently, Stein *et al*. (2017) collected *A. halleri* leaves from 1466 plants from soils classified as non-metalliferous sites based on a multifactorial metal analysis rather than the strict thresholds defined in Bert *et al*. (2002). Due to their extensive sampling, Stein *et al*. (2017) found much broader ranges of heavy metal concentrations in soils and plants from non-metalliferous sites than earlier studies of *A. halleri*, with some plants accumulating more than 50,000 µg g^-1^ of Zn in non-contaminated sites. This is characteristic of a deficiency response for hyperaccumulating species (Hanikenne *et al*. 2008; Krämer 2010) where *A. halleri* became very efficient at extracting large amounts of Zn from soils when Zn concentrations in the soils are low. Similarly, we found that *A. kamchatica* plants from non-modified habitats where Zn concentrations were below 100 µg g^-1^, such as four populations from Alaska, USA (see Figure 1), had up to 2,000 µg g^-1^ of Zn in leaves. This indicates that *A. kamchatica* plants exposed to low Zn concentrations in native, non-modified soils can still accumulate substantial amounts of Zn. This is consistent with constitutive hyperaccumulation in *A. kamchatica* as in *A. halleri*.

A fortunate byproduct of our sampling revealed that some sites containing *A. kamchatica* showed evidence of serpentine soils (those with high Mg and Ni and low Ca:Mg ratio), particularly those at high elevation (>3,000 m) in the Northern Alps of Central Japan. These serpentine soils have been previously documented, but surveys of plants growing on these soils did not report the presence of *A. kamchatica* (Hatano and Matsuzawa 2008) or other synonyms. The elevated levels of Mg and Ni in plants down-stream from these sites at the North and South River Matsu sites suggests runoff may carry these ions to lower elevations. Adaptations to serpentine soils have been identified in North American and British populations of *A. lyrata* (Turner *et al*. 2010; Arnold *et al*. 2016), one of the diploid parents of *A. kamchatica*. It is therefore possible that *A. kamchatica* may be adapted to transport heavy metals such as Cd and Zn derived from *A. halleri*, and have tolerance to serpentine soils derived from *A. lyrata*. The diversity of soil types in Japan where *A. kamchatica* is found suggests that the ability to inhabit highly variable environments may be due to parental merging of stress tolerance in allopolyploid species (Godfree *et al*. 2017; Van de Peer *et al*. 2017).

### Experimental treatments show zinc accumulation is a constitutive trait in A. kamchatica

Growing naturally collected *A. kamchatica* plants in hydroponic conditions allowed us to estimate quantitative genetic variation in Zn accumulation by removing the environmental heterogeneity found in the natural populations. Moreover, the hydroponic conditions allowed us to quantify Zn in roots which would be extremely difficult in naturally collected samples. Our experiments demonstrated an 8-fold difference between the lowest and highest genotypes for Zn accumulation in leaf tissues, indicating significant variation for Zn hyperaccumulation in *A. kamchatica*. When plants were treated with the same Zn treatment, the average Zn accumulation in leaves among the *A. kamchatica* genotypes exceeded the threshold defining hyperaccumulating species (> 3,000 µg g^-1^) (Krämer 2010). Despite a large amount of standing genetic variation for Zn hyperaccumulation in the species, *A. kamchatica* clearly has constitutive hyperaccumulation ability.

Further support for constitutive Zn hyperaccumulation is found by the lack of a significant relationship between Zn concentrations in native soils with leaf accumulation levels using *A. kamchatica* germplasm from where soils were collected. Similarly, even with large sampling of plant genotypes of *A. halleri* from non-metalliferous sites, no significant correlation was found between Zn concentrations in native soils and plants from these sites that were grown experimentally using Zn amended soils (see Fig. 3c in Stein *et al*. 2017). Consistent with our study, Stein *et al*. (2017) found that the largest proportion of variance in Zn accumulation in plants was explained by plant genotype and not soil concentrations, and that most of the variance was detected in plants that came from sites with < 200 µg g^-1^ of Zn in soils. In addition to our experimental results, constitutive hyperaccumulation of Zn is also supported by our finding of plant tissues collected from natural populations that had large amounts of Zn, even when Zn concentrations were low in the soils where plant tissues were collected.

Our experimental results for *A. kamchatica* are consistent with what has been shown for *A. halleri* in several studies using hydroponic Zn treatments (Bert *et al*. 2002; Talke 2006; Hanikenne *et al*. 2013) and strongly supports that Zn accumulation was inherited from *A. halleri*. We were able to directly test inheritance of hyperaccumulation by using a synthesized allopolyploid generated from *A. halleri* and *A. lyrata*. The synthesized *A. kamchatica* had 73% of accumulation of Zn in the leaves compared with the parental *A. halleri* strain (a reduction or attenuation of 27%) used in the same experiment (Figure 3, Table S5a). This is considerably higher than most natural genotypes which have on average about 0.5 of the accumulation of Zn in leaves compared to *A. halleri*. Moreover, accumulation of Zn in leaves of the synthetic polyploid was orders of magnitude higher than that of the *A. lyrata* parent (see Fig. 1 in Paape *et al*. 2016), which clearly demonstrated that hyperaccumulation can be retained following hybridization between the divergent parental species, but that the *A. lyrata* genome has weakening or attenuating effects on the trait.

### Larger quantities of Zn retained in the roots of A. kamchatica compared with A. halleri

The hydroponic experiments using static conditions showed that *A. kamchatica* accumulated substantial amounts of Zn when given a large treatment of Zn (500 or 1000 µM) for one week, and some genotypes accumulated as much as *A. halleri*. However, the long duration of the treatment (one week) may result in Zn transport approaching equilibrium levels in *A. kamchatica,* where given enough time, the polyploid could potentially accumulate levels of Zn similar to *A. halleri*. Testing Zn accumulation over a gradient of treatments for a short period of time provides a better picture of the relative efficiency by which *A. kamchatica* and *A. halleri* can physiologically transport Zn from roots to shoots. At all treatment conditions, *A. halleri* accumulated significantly more Zn in the leaves than *A. kamchatica*. While *A. kamchatica* accumulated up to 1,000 and 3,000 µg of Zn in leaves at the 100 µM and 1,000 µM treatments respectively, more Zn was retained in the roots compared with *A. halleri*. The shoot to root ratio of Zn concentration was greater than or equal to one for the *A. halleri* genotype used in our experiment, which is a similar result to a previous study that tested Zn accumulation over a gradient of treatments using a European *A. halleri* genotype (Talke 2006). This demonstrated efficient root to shoot transport at low, intermediate, and high treatments for *A. halleri*.

By contrast, the shoot to root ratio of Zn concentration for *A. kamchatica* was equal to or less and one-half for the three genotypes used in our gradient experiment. Notably, the highest shoot to root ratio was at the control condition (1 µM) for both *A. halleri* and *A. kamchatica*, which is consistent with a Zn deficiency response due to constitutive expression of heavy metal transporters (Talke 2006; Hanikenne *et al*. 2008). The hydroponic treatments demonstrated experimentally how plants can accumulate high levels of Zn when exposed to both low and high levels of Zn, which is important for understanding the relationship between soil concentrations and accumulation by plants in natural conditions.

We suggest two reasons why an allopolyploid derived from two parents that were divergent for heavy metal hyperaccumulation would show a reduction in the hyperaccumulation trait compared to the diploid hyperaccumulator parent (*A. halleri*). First, a reduction or attenuation compared to the diploid parents in total expression levels of metal transporters is expected due to allopolyploidization. A reduction by about half of the expression was detected in several important heavy metal transporters in the *A. halleri*-derived homeologs in *A. kamchatica* compared with the orthologous genes in *A. halleri* using RNA-seq data (Paape *et al*. 2016). Half of the expression levels of metal transporters is consistent with the roughly half of the level of Zn accumulation in leaves of natural accessions of *A. kamchatica* compared with *A. halleri*. In other words, allopolyploidization results in a state of fixed heterozygosity of functionally duplicated gene copies (homeologs), reducing expression of homeologs compared to the diploid parents. Fixed heterozygosity can result in a trait that resembles a balanced polymorphism with a semi-intermediate phenotype. It is important to note that while the hyperaccumulation trait is reduced compared to *A. halleri* by about one-half, it is an order of magnitude or greater than the non-hyperaccumulating parent *A. lyrata* (Paape *et al*. 2016). Second, there are likely inhibiting genetic factors derived from the *A. lyrata* parental genome. Here it is expected that these genetic factors act to prevent the transport of toxic heavy metals to leaf tissues to limit toxicity in leaves. In this case, there is genomic antagonism resulting from the divergent parental genomes.

### Expression bias in A. halleri derived homeologs

Because Zn hyperaccumulation was likely inherited from *A. halleri*, we expected that homeolog expression ratios would show a pattern consistent with a parental legacy effect of gene expression (Buggs *et al*. 2014). Using pyrosequencing assays we found expression ratios of homeologs for six genes with roles in heavy metal hyperaccumulation or metal tolerance each had higher expression of the H-origin copy. This suggests homeolog specific expression is maintained by *cis*-regulatory differences (Shi *et al*. 2012; Yoo *et al*. 2014). In addition to *cis*-regulation, the major genes known to be responsible for root to shoot transport of Zn (*HMA4*) and detoxification of tissues by vacuolar sequestration (*MTP1*) in *A. halleri*, have high expression in *A. halleri* because they are duplicated in *A. halleri* (3 copies for *HMA4*, Hanikenne *et al*. 2008; 3 - 5 copies for *MTP1* Shahzad *et al*. 2010) while they are single copy in the non-hyperaccumulationg congeners *A. lyrata* and *A. thaliana* (Briskine *et al*. 2017). Therefore, these two genes show significantly higher expression when compared with orthologs from non-hyperaccumulating congeners (Becher *et al*. 2004; Filatov *et al*. 2006; Paape *et al*. 2016) due to increased copy number. The gene *HMA3* also has a putative role in vacuolar sequestration of Zn (Morel *et al*. 2008), similar to *MTP1,* but is only found in a single copy in both *A. halleri* and *A. lyrata* (Paape *et al*. 2018). This gene also has very high allele specific expression in *A. halleri* (Talke 2006) and strong H-origin bias in *A. kamchatica*, which must be due to *cis*-regulatory differences. Together, these three genes (*HMA3, HMA4, MTP1*) had the strongest bias in H-origin homeolog expression in both leaves and roots, showed the least amount of variation in homeolog expression ratios among the 15 genotypes tested, and had stable expression ratios before and after Zn treatment. These features are consistent with constitutive gene expression that was inherited (Buggs *et al*. 2014) from the hyperaccumulating diploid parent *A. halleri*, which would be essential for retaining the hyperaccumulation phenotype in the species-wide collection of *A. kamchatica* examined in this study.

The *IRT3* and *NRAMP3* homeologs also showed H-origin expression bias consistent with previous differential expression studies comparing *A. halleri* with *A. lyrata* or *A. thaliana* in Zn treatment studies (Filatov *et al*. 2006; Talke 2006), but their direct role in Zn hyperaccumulation is less clear (Thomine *et al*. 2003). Moreover, both *IRT3* and *NRAMP3* showed much larger variation in expression ratios than *HMA4* or *MTP1* which may reflect greater constraint on constitutive expression of the latter two genes.

Most importantly, Zn treatment had no significant effect on the expression ratio of five of the six genes, demonstrating constitutive expression for genes with known or putative roles in Zn hyperaccumulation. The gene *MTP3* was an exception and showed a significant change in homeolog ratios in the root tissues of many genotypes following the Zn treatment. The ratio change was most likely the result of up-regulation of the L-origin homeolog in roots, which was previously shown in one of the genotypes used in the current study using RNA-seq (Paape *et al*. 2016). It has been shown that *MTP3* has a role in preventing heavy metal transport to the shoots by sequestering Zn in the vacuoles in roots in *A. thaliana* (Arrivault *et al*. 2006). Because we assume this gene would have a similar role in *A. lyrata*, it is a potential *A. lyrata*-derived inhibiting factor that would contribute to reduced leaf hyperaccumulation in *A. kamchatica*.

## Conclusions

Adaptability and range expansion in polyploids has been discussed for several decades (Stebbins 1971; Soltis *et al*. 2014; Van de Peer *et al*. 2017) but empirical examples of quantitative traits that have ecological relevance have been lacking (Godfree *et al*. 2017). The ability of allopolyploid species to merge parentally inherited adaptations to climate or soils has been proposed to confer greater plasticity compared with the diploid progenitors (Yoo *et al*. 2014; Shimizu-Inatsugi *et al*. 2017), which could be a mechanism to mediate niche expansion (Blaine Marchant *et al*. 2016). We have shown that an allopolyploid species with a broad habitat has retained heavy metal hyperaccumulation from one of the diploid parents as evidenced by field and experimental analyses of species wide population samples. Zinc hyperaccumulation appears to be a constitutive trait in *A. kamchatica*, driven by parentally inherited gene expression of important genes involved in the trait. We also discovered significant quantitative variation for Zn accumulation in the allopolyploid species which would further increase adaptability to broader and more diverse environments and soil types (Barrett & Schluter 2008; Matuszewski *et al*. 2015).

We suggest that the inheritance of hyperaccumulation from *A. halleri* conferred advantages instantaneously at the polyploid speciation, estimated to be approximately 100,000 years ago (Paape *et al*. 2018) by the following scenario. First, *A. kamchatica* became tolerant to soils with toxic levels of heavy metals that were present in natural populations due to geological processes. Then, during the past few thousand years, soils became contaminated by human activities, and the tolerance worked as a pre-adaptation for modified environments (a similar scenario has been proposed for *A. halleri*, Meyer *et al*. 2016). In contrast to *A. halleri*, *A. kamchatica* was not found in extremely contaminated sites such as mines, consistent with attenuated Zn hyperaccumulation. This would have contributed to a distinct, intermediate habitat and species distribution of *A. kamchatica* compared with the diploid parents. Second, Zn concentration in the leaves of the majority of the natural *A. kamchatica* populations were above 1,000 µg g^-1^. This level was shown to be effective for insect defense in *A. halleri* (Kazemi-Dinan *et al*. 2014). In addition, we found *A. kamchatica* living on serpentine soils, which has also been found in the other diploid parent *A. lyrata* (Turner *et al*. 2010; Arnold *et al*. 2016). We hypothesize that *A. kamchatica* expanded its habitats by combining heavy metal and serpentine tolerance from *A. halleri* and *A. lyrata*. Our study represents a promising example in which the inheritance of genetic toolkits for soil adaptations likely contributed to the habitat expansion of an allopolyploid species. New genomic and transcriptomic capabilities in *A. kamchatica* combined with functional genetics (Yew *et al*. 2017) and self-compatibility (Tsuchimatsu *et al*. 2012) provides a unique opportunity to study the genetics of edaphic and climatic adaptation of a polyploid species (Shimizu *et al*. 2011).

## Methods

### Plant and soil material from natural populations

*Arabidopsis kamchatica* (Fisch. ex DC.) K. Shimizu & Kudoh (Shimizu *et al*. 2005) is an allotetraploid species distributed in East Asia and North America. The diploid parental species *A. halleri* and *A. lyrata* each possess 8 chromosomes (*n* = 8, 2*n* = 2*x* = 16) and the allopolyploid hybrid has 4*n* = 4*x* = 32 chromosomes. Leaf tissues were collected from 40 *A. kamchatica* populations in Japan, Russia and Alaska, USA to quantify heavy metals in natural conditions (Tables S1) using inductively coupled mass spectrometry (ICP-MS). For most populations we collected leaf tissues from at least three plants. We reported values for leaf accumulation based on the mean and median of the replicates at each site, while often individual plants greatly exceeded the mean for any population. Soil samples were collected from the majority of these sites to measure metal ion concentrations (Table S2). Because many of the sites where *A. kamchatica* was collected showed clear signs of human modification, we grouped the sites based on: 1) Modified populations are on or next to paved roads (which includes concrete and/or asphalt, paved slope or paved riverside), or next to the base of a building. 2) Natural populations are on native soil, free from concrete, asphalt, and in-native dirt. We also collected leaf tissues from *A. halleri* subsp. *gemmifera* from two sites in Japan and two sites in Russia. Soil samples were collected from the two Japanese sites, which are known mine sites (Tada mine (TADA) and Omoidegawa (OMD), Japan, Briskine *et al*. 2017). The Russian samples were taken from herbarium specimens and soils from these locations were not collected.

### Hydroponic plant growth experiments

We conducted hydroponic experiments to measure Zn accumulation in leaves and roots using natural genotypes of *A. kamchatica* and *A. halleri* from germplasm collected from natural populations (Table S5). Seeds of *A. kamchatica* were germinated on phytoagar (0.8%) and a mixture of oligonutrients (25 µm H_3_BO_3_, 5 µm MnCl_2_, 1 µm ZnSO_4_, 0.5 µm CuSO_4_, 50 µm KCl, and 0.1 µm Na_2_MoO_4_) plated on a square 8 cm x 8 cm petri dish until the seeds started to germinate (about one week). We used 1000 µL pipet tip boxes (ca. 700 mL volume) for the hydroponic chambers so that seedlings could be grown in 0.5 mL thermo-PCR tubes using the 96-well insert (about 20 seedlings per box for adequate spacing). The seedlings were then transplanted in 0.5 mL thermo-PCR tubes also filled with phytoagar solution and placed the pipet tip boxes. The hydroponic solution was prepared according to Paape *et al*. (2016) which was composed of: 4 mM KNO_3_, 1.2 mM Ca(NO_3_)_2_, 0.8 mM MgSO_4_, 0.8 mM KH_2_PO_4_, 0.8 mM NH_4_Cl, and 5 μM Fe(III)EDTA. A separate 1 L stock of oligoelements was made with the following elements and 16.25 mL of oligonutrients was added to the final 5 L solution: 0.2 mM KCL, 0.12 mM H_3_BO_3_, 0.04 mM MnSO_4_, 4 μM CuSO_4_, 1 μM ZnSO_4_, and 1 μM (NH_4_)_6_Mo7O_2_. Ten-liter batches were mixed in one container and then dispensed to individual hydroponic containers. The final pH was adjusted to 5.6–5.8.

To keep the moisture suitable for the immature seedlings to grow, a plastic bag was wrapped around each container to maintain high humidity. The container was placed near natural light for the duration of 3-4 days. The boxes were then moved into a growth chamber (16 hrs. light, 8 hrs. dark at 20°C) and the plastic bag removed. The boxes containing the seedlings were placed on a 40 cm x 60 cm green tray (4 boxes per tray) containing 1 cm of water, and covered with a plastic lid in order to maintain high humidity. After the roots grew to about 0.5 cm, the bottom of the plastic tube was cut with scissors, allowing the root to elongate into the box containing hydroponic solution. A similar procedure was used for *A. halleri* by placing freshly cut clones (i.e. ramets from a living plant) directly into the 0.5 mL tubes. Once the plants achieved the three to four leaf stage, the lid was removed to allow direct light. Zn supplements (treatments using 1 – 1000 µM ZnSO_4_ supplements) were added after about 4.5 weeks of plant growth (with slight variability among genotypes because of seed germination).

We conducted three Zn treatment experiments. The first experiment used a supplement of 500 µM Zn added to the hydroponic solution for a period of 7 days. This experiment included 19 natural *A. kamchatica* genotypes and one synthetic polyploid that was generated from *A. halleri* subsp. *gemmifera* (w302) and *A. lyrata* subsp. *petraea* (Table S1). We also used two naturally collected *A. halleri* genotypes that we maintain in the lab, TADA (*A. halleri* subsp. *gemmifera*, originally collected from the Tada Mine, Japan, w302 strain ID) and BOD (*A. halleri* subsp. *halleri* collected from Boden, Switzerland, w875 strain ID) which can be clonally propagated for experiments. We began with 10 – 15 replicates per plant genotype depending on germination. Plants that initially germinated but died in the immature stage were discarded from the experiment. The final number of biological replicates for each accession varied from 5 – 13. Each box contained a single plant genotype to avoid root contamination between genotypes during sampling and harvesting. The boxes were randomly placed on 40 cm x 60 cm green trays and their positions were regularly rearranged throughout the growth period.

Leaves and roots were harvested from plants grown after 5.5 weeks, with the final week of exposure to zinc treatment as described above. The second experiment included 9 *A. kamchatica* genotypes with a supplement of 1000 µM Zn added to the hydroponic solution for a period of 7 days. The third experiment (“Zn gradient experiment”) varied Zn concentrations at ten-fold increments of Zn treatments, 1 µM (control condition), 10 µM, 100 µM and 1000 µM, to compare relative accumulation at different concentrations and between species and genotypes. Three genotypes of *A. kamchatica* (MUR, SAK, PAK, 10 – 12 replicates each) and one *A. halleri* (TADA, 6 replicates) were used for this experiment. Plants were grown for ca. 4.5 weeks prior to the treatments. Leaves and roots were harvested after 48 hours of exposure to each of the four treatments to measure short term uptake of Zn. For tissue collection for metal analysis in in all three experiments, leaves and roots (approximately 2–5 mg dry weight) were harvested from plants following Zn treatments as described above. During harvesting, root tissues were washed in 150 mL of cold solution of 5 mM CaCl_2_ and 1mM MES-KOH (pH 5.7) for 30 min, followed by a wash in 150 mL cold water for 3 min. The tissues were then collected in paper envelopes and dried at 60°C for 2 days. Root tissues were rinsed again with 18 MΩ water and placed into Pyrex digestion tubes. Leaf and root tissues were dried at room temperature and then at 50°C for 24 h.

### Elemental analysis in plant tissues and soil samples

Measurements of heavy metal concentrations were performed as described by Lahner *et al*. (2003) at the Institute of Terrestrial Ecosystems at ETH Zurich and by Paape et al. (2016). Plant tissues were weighed and placed into 50 ml tubes. Samples were placed into an oven at 92°C to dry for 20 hours prior to ion measurement. After cooling, reference samples for each ion were weighed. Samples were digested in a microwave oven with 2 mL concentrated nitric acid (HNO_3_, ACS reagent; Sigma-Aldrich) and 30% hydrogen peroxide (Normapur; VWR Prolabo) and diluted to 10 mL with 18 MΩ water. Analytical blanks and standard reference material (WEPAL IPE 980) were digested together with plant samples. ICP-MS was used for elemental analysis with samples and reference standards. To correct for instrumental drift, an internal standard with yttrium and indium were added to the samples. All samples were normalized to calculate weights, as determined with a heuristic algorithm using the best-measured elements, the weights of the samples, and the elemental solution concentrations. Soil samples were weighed and finely ground, then dried at 40°C. Soils were digested using the DigiPREP MS digestion system (SCP Science, Canada) and 2 M HNO_3_ for 90 min at 120°C. The samples were then cooled and diluted with up to 50 ml of nanopure water. The digested soils were then filtered using 41 Whatman filter paper into 50 ml centrifuge tubes. Samples were then diluted for measurement using inductively coupled plasma optical emission spectrometry (ICP-OES) at ETH Zurich.

### RNA extraction and Pyrosequencing

Leaf and root tissues from 15 *A. kamchatica* genotypes were harvested from three replicates grown in hydroponic solution at time zero (control) and at 48 hours following the replacement of the hydroponic solution containing a 500 µM supplement of Zn (from the 500 µM Zn experiment described above). Leaf and root tissues were flash frozen in liquid nitrogen and stored at -80°C. RNA was extracted with TRIzol (Invitrogen) and purified with the RNeasy Mini Kit (Qiagen). RNA concentrations were measured using Nanodrop (Thermo Scientific). The RNA samples were reverse transcribed to cDNA using the High Capacity RNA-to-cDNA kit (Invitrogen).

To measure homeolog expression ratios of metal-ion transporter genes, we used pyrosequencing on reverse transcribed cDNA templates. Pyrosequencing is a PCR based method that can detect the relative abundance of homeolog-specific single-nucleotide polymorphisms (SNPs) which will vary in abundance based on homeolog expression levels (Akama *et al*. 2014; Paape *et al*. 2016). Based on previous studies we selected the orthologous *A. thaliana* genes *HMA3* (AT4G30120), *HMA4* (AT2G19110), *MTP1* (AT2G46800), *MTP3* (AT3G58810), *NRAMP3* (AT2G23150) and *IRT3* (AT1G60960) (Table S8). We used coding sequence alignments in fasta format, generated from resequencing data of *A. kamchatica* homeologs (Paape *et al*. 2018). In *A. halleri*, the gene *HMA4* is a tandemly triplicated gene and *MTP1* has at least three copies (Hanikenne *et al*. 2008, 2013; Shahzad *et al*. 2010; Briskine *et al*. 2017), but only one of the three copies from our reference genome for either of these genes was used for assay design as polymorphism between the copies is low. These two genes are present in only a single copy in *A. lyrata* (Briskine *et al*. 2017; Paape *et al*. 2018), the other diploid parent of *A. kamchatica*.

To design PCR primers, sequencing primers, and SNP assays for each gene, we used the PyroMark Assay Design v2.0 software (Qiagen) at the Genomic Diversity Center (GDC), ETH, Zurich. To design pyrosequencing assays, we aligned coding sequences of homeologous gene copies for each of the six genes above to detect SNPs between *A. halleri* (H-origin) and *A. lyrata* (L-origin) derived copies. We searched for target SNPs between two conserved primers that contained 2 – 3 target SNPs in regions less than 200 bp using the PyroMark software (Qiagen). The amplified PCR fragments were sequenced using the PyroMark ID software (Qiagen) at CDC, ETH, Zurich. The amplification peaks were analyzed using the PyroMark software using the AQ (allele quantification) mode to determine the SNP amplification ratio. The ratios of the 2 – 3 target SNPs obtained for each gene fragment were averaged to estimate the H and L – origin homeolog expression ratios. Standard deviations for each gene were estimated from the three biological replicates. For the H-origin homeologs of *HMA4* and *MTP1*, we assumed the H-origin expression is the sum of the three duplicated copies derived from *A. halleri*.

## Statistical analysis

For the leaf and soils collected from natural populations we calculated the mean, median, range and standard deviations for each population which typically consisted of three replicates. To determine whether there was a correlation between leaf accumulation of Cd and Zn and soil concentrations of these heavy metals, we used Pearson’s correlation coefficients. We used ANOVA and a generalized linear model (glm) to detect quantitative variation among genotypes of *A. kamchatica* in Zn accumulation in leaf tissues, root tissues or leaf to root ratios in the 500 µM Zn experiment. We assessed whether leaf Zn, root Zn, and leaf/root ratio of Zn vary among genotypes of *A. kamchatica*. Each of the three response variables were analyzed separately as the focus was on variation within tissue and not on Zn and tissue interactions. We performed the analysis with and without the outlier genotype, MAG to determine whether significant trait variation was influenced by this genotype. Due to unbalance, heteroscedasticity, and non-normality, the models did not satisfy the assumptions of linear models. The effect of genotype was therefore examined using a glm with a Gamma distribution (for continuous response variable) with inverse link function. For the 10-fold gradient experiment we compared means and standard deviations of three replicates of each genotype at each Zn treatment condition. Here we were primarily concerned with comparisons between *A. halleri* (the diploid parent) and *A. kamchatica*, and secondly whether the three *A. kamchatica* genotypes used in the experiment showed significant differences at each condition. Significant differences were evaluated by non-overlapping means or standard deviations.

We used linear models to determine the effects of genotype, gene (function), treatment, and tissue on homeolog expression ratios (estimated using pyrosequencing). The expression ratios vary from 0 to 1 according to the relative expression of either homeolog. If there is equal expression of both homeologs, H-origin = 0.5 and L-origin = 0.5. Therefore, we used the H-origin copy expression as the dependent variable. We used the following linear model formula that included both tissue types together in the model: H-origin ratio ∼ genotype + gene + treatment + tissue + genotype x gene (interaction term) + gene x treatment (interaction term) + gene x tissue (interaction term) + gene x treatment x tissue (interaction term). The significance of each explanatory variable was summarized by sum of squares and *F*-statistics in an ANOVA table (See Table S9). We then separated the datasets into leaf or root only and repeated the linear model procedure on both datasets separately, using the following model: H-origin ratio ∼ gene + treatment + genotype + gene x treatment (interaction term) + gene x genotype (interaction term). All analyses were performed in software R, version 3.3.3.

## Acknowledgments

We thank Rainer Schulin and Bjoern Studer at the Institute of Terrestrial Ecosystems at ETH Zurich for the metal ion analysis facilities as well as technical support and useful discussions. Enrico Martinoia formerly at UZH Plant Biology for material contributions to hydroponics experiments and valuable discussions about experimental design and results. The Genetic Diversity Center at (GDC) for software and equipment to perform pyroseqencing. Part of field leaf and soil samples were provided by Miss Hinako Kanai. The study was supported by the European Union’s Seventh Framework Programme for research Swiss Plant Fellows to TP, the University Research Priority Program of Evolution in Action of the University of Zurich to TP and KKS, a JST CREST grant (number JPMJCR16O3) to K.K.S. and T.K., Swiss National Science Foundation to K.K.S., MEXT KAKENHI Grant Numbers 16H06469 and 18H04785 to KKS, the Special Coordination Funds for Promoting Science and Technology from the MEXT, Japan to TK, an Inamori Foundation research grant to TK, and JSPS Grant-in-Aid for Scientific Research (2277023) to TK.

## Author contributions

TP and KKS conceived of the study. TP and TC designed and performed experiments and prepared samples for analysis. TP designed pyrosequencing assays and TC performed pyrosequencing. TP, YO, AH, TK and KKS collected field samples. TP and TC constructed datasets. TP and RA performed statistical analyses. TP drafted the manuscript. TP, RA, TK and KKS revised the final draft.

## Supplementary Figures

**Figure S1.**
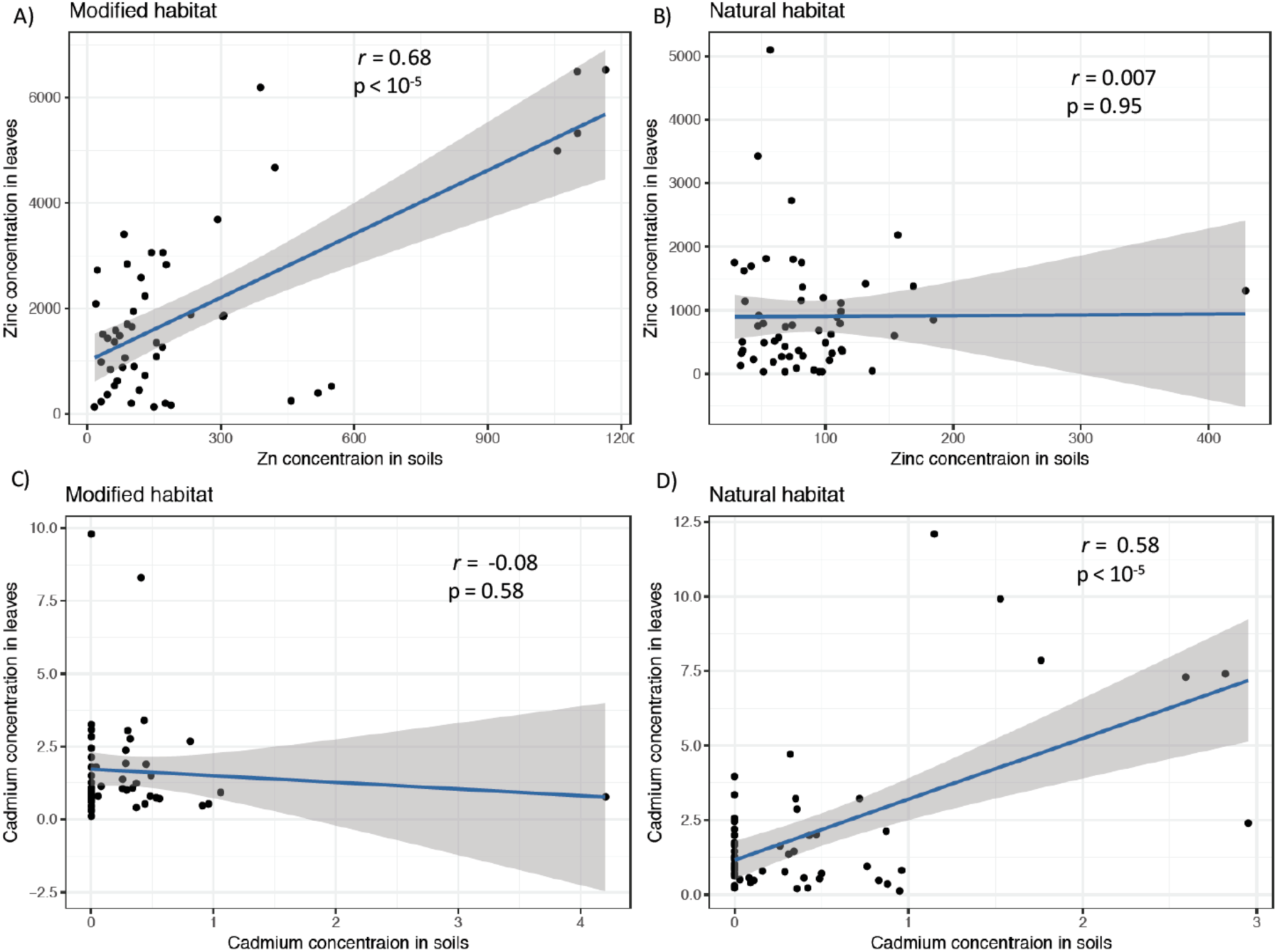
Correlation arrays of leaf accumulation of cadmium (Cd) (left) and zinc (Zn) (right) with corresponding soil concentrations of the two heavy metals respectively.

**Figure S2.**
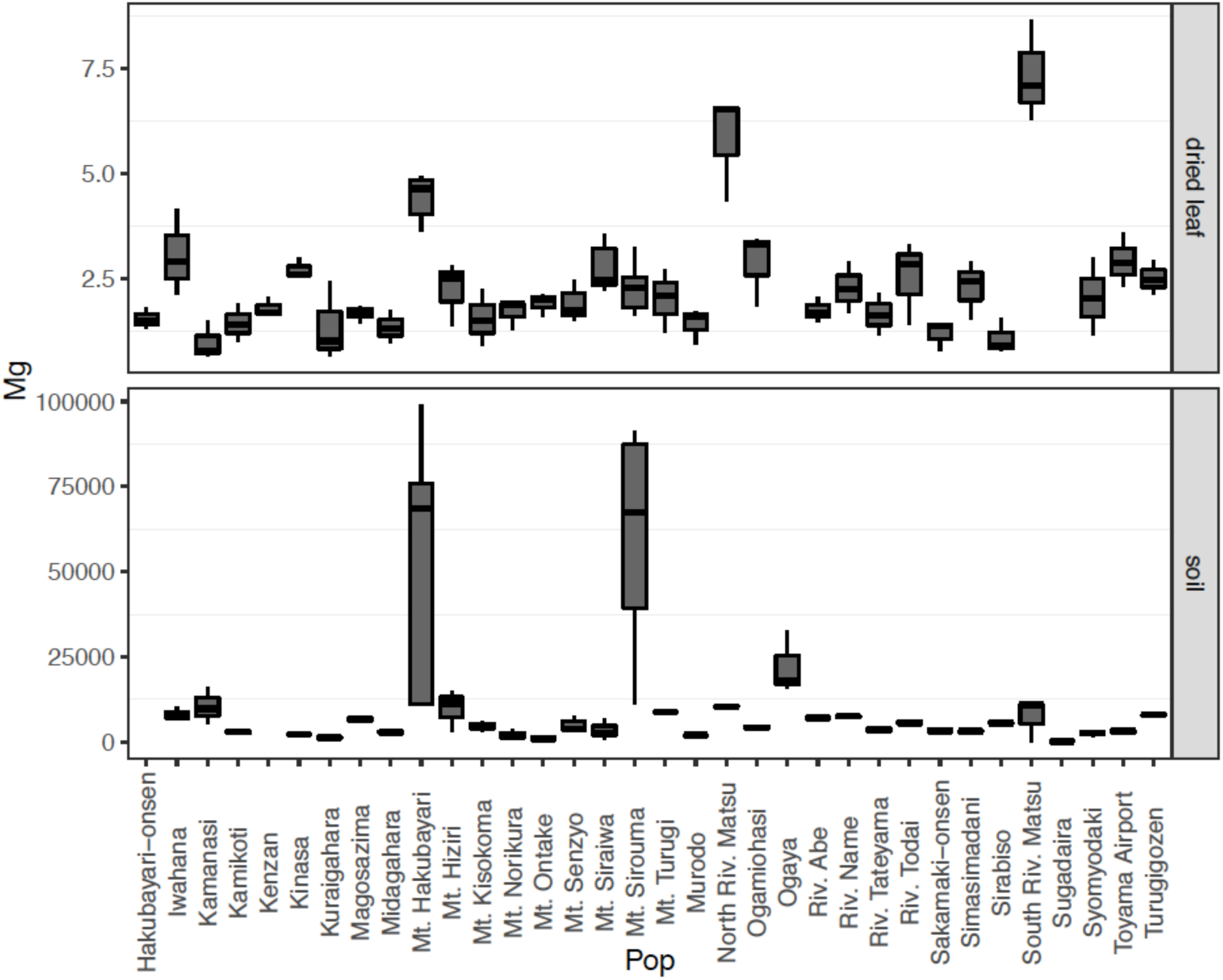
Magnesium (Mg) concentrations in leaf tissues (upper panel) and Mg concentrations in soils at each collection site. Boxplots show center line: median; box limits: upper and lower quartiles; whiskers: 1.5 × interquartile range; points: outliers.

**Figure S3.**
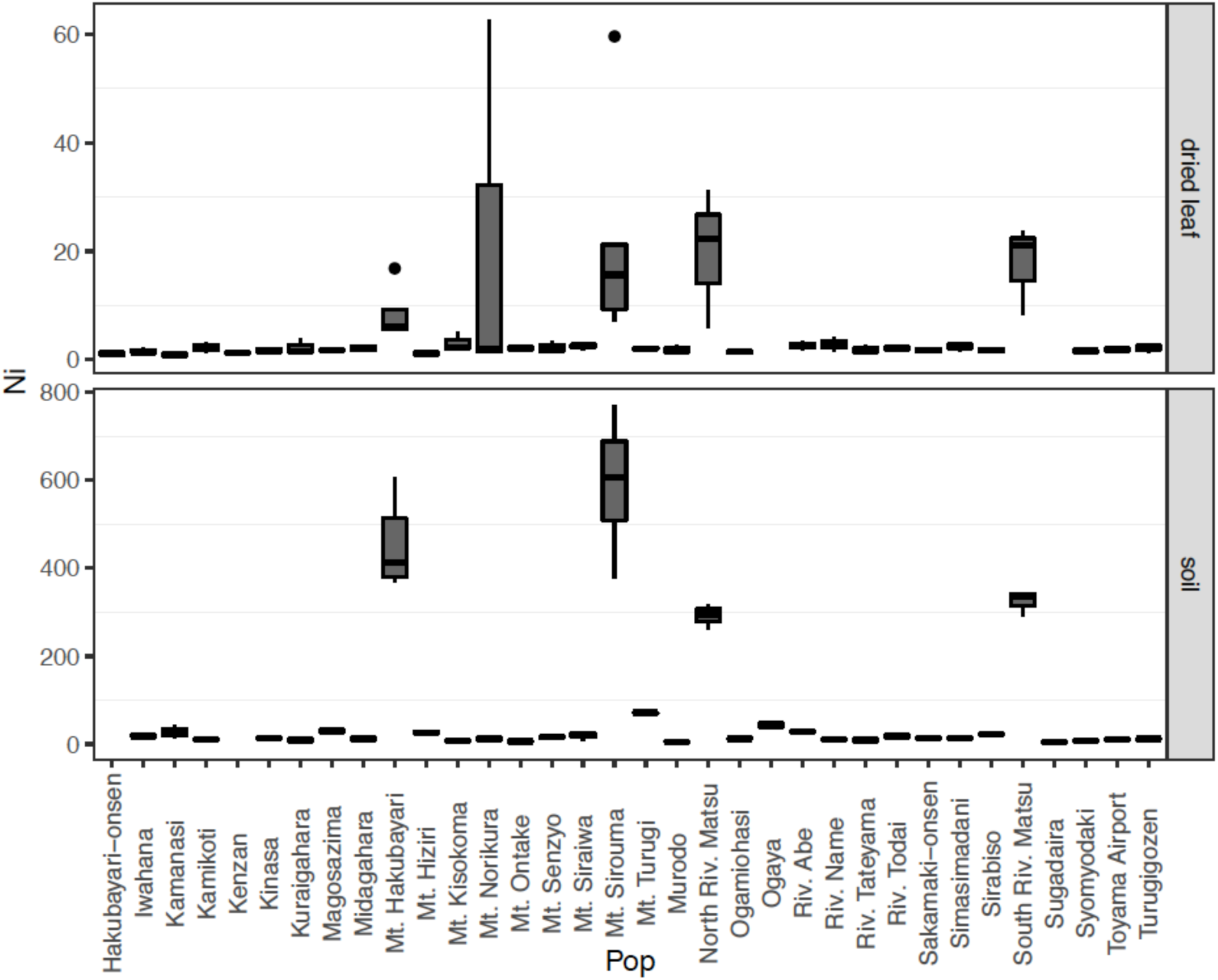
Nickel (Ni) concentrations in leaf tissues (upper panel) and Ni concentrations in soils at each collection site. Boxplots show center line: median; box limits: upper and lower quartiles; whiskers: 1.5 × interquartile range; points: outliers.

**Figure S4.**
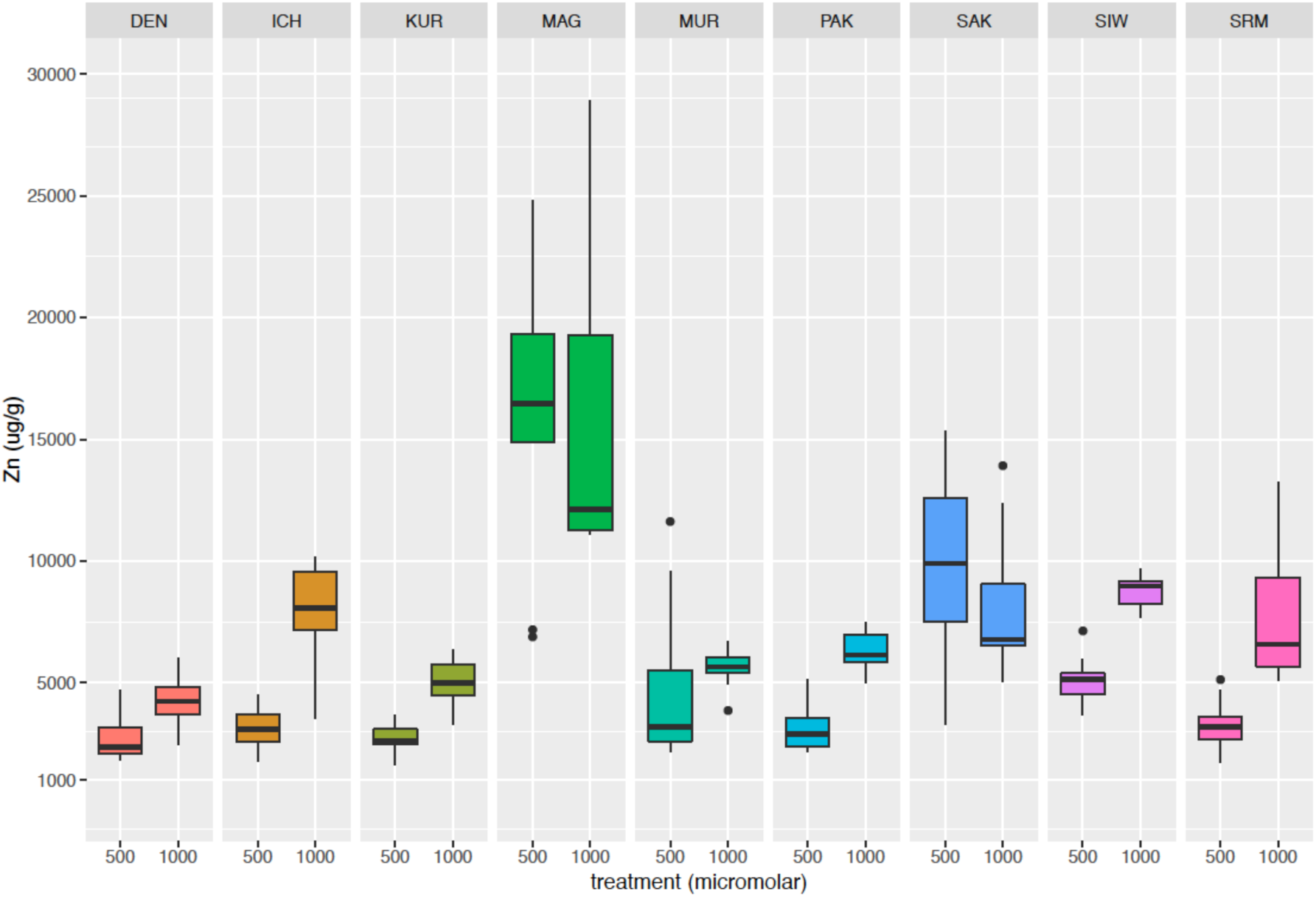
Zn accumulation in leaves of nine *A. kamchatica* accessions following 7 days of Zn treatments of 500 and 1000 µM of Zn in two independent experiments (the same plants were not exposed to both 500 and 1000 µM treatment conditions but were grown in separate hydroponic containers in either treatment). Significant increases in mean Zn accumulation were observed in the 1000 µM treatment compared with the 500 µM treatment for all except the MAG and SAK genotypes. The MAG and SAK genotypes showed no significant increase in mean Zn accumulation between 500 and 1000 µM treatments.

